# Exploiting single-cell quantitative data to map genetic variants having probabilistic effects

**DOI:** 10.1101/040113

**Authors:** Florent Chuffart, Magali Richard, Daniel Jost, Hélène Duplus-Bottin, Yoshikazu Ohya, Gaël Yvert

**Affiliations:** Univ Lyon, ENS de Lyon, Univ Claude Bernard, CNRS UMR 5239, INSERM U1210, Laboratory of Biology and Modelling of the Cell, 46 allée d’Italie Site Jacques Monod, F-69007, Lyon, France; Univ Lyon, ENS de Lyon, Univ Claude Bernard, CNRS, Laboratory of Physics, 46 allée d’Italie Site Jacques Monod, F-69007, Lyon, France; University Grenoble Alpes, CNRS, TIMC-IMAG lab, UMR 5525, Grenoble, France.; Department of Integrated Biosciences; Graduate School of Frontier Sciences; University of Tokyo; Kashiwa, Chiba; Japan.

**Keywords:** Quantitative Genetics, Missing Heritability, QTL, GWAS, incomplete penetrance, small-effect variants, yeast, galactose, GAL, single-cell, gene expression noise, stochasticity

## Abstract

Despite the recent progress in sequencing technologies, genome-wide association studies (GWAS) remain limited by a statistical-power issue: many polymorphisms contribute little to common trait variation and therefore escape detection. The small contribution sometimes corresponds to incomplete penetrance, which may result from probabilistic effects on molecular regulations. In such cases, genetic mapping may benefit from the wealth of data produced by single-cell technologies. We present here the development of a novel genetic mapping method that allows to scan genomes for single-cell Probabilistic Trait Loci that modify the statistical properties of cellular-level quantitative traits. Phenotypic values are acquired on thousands of individual cells, and genetic association is obtained from a multivariate analysis of a matrix of Kantorovich distances. No prior assumption is required on the mode of action of the genetic loci involved and, by exploiting all single-cell values, the method can reveal non-deterministic effects. Using both simulations and yeast experimental datasets, we show that it can detect linkages that are missed by classical genetic mapping. A probabilistic effect of a single SNP on cell shape was detected and validated. The method also detected a novel locus associated with elevated gene expression noise of the yeast galactose regulon. Our results illustrate how single-cell technologies can be exploited to improve the genetic dissection of certain common traits.

**AUTHOR SUMMARY:** Genetic association studies are usually conducted on phenotypes measured at the scale of whole tissues or individuals, and not at the scale of individual cells. However, some common traits, such as cancer, can result from a minority of cells that adopted a special behavior. From one individual to another, DNA variants can modify the frequency of such cellular behaviors. The body of one of the individuals then harbours more misbehaving cells and is therefore predisposed to a macroscopic phenotypic change, such as disease. Such genetic effects are probabilistic, they contribute little to trait variation at the macroscopic level and therefore largely escape detection in classical studies. We have developed a novel statistical method that uses single-cell measurements to detect variants of the genome that have non-deterministic effects on cellular traits. The approach is based on a comparison of distributions of single-cell traits. We applied it to colonies of yeast cells and showed that it can detect mutations that change cellular morphology or molecular regulations in a probabilistic manner. This opens the way to study multicellular organisms from a novel angle, by exploiting single-cell technologies to detect genetic variants that predispose to certain diseases or common traits.

## INTRODUCTION

The aim of modern genetics is to identify DNA variants contributing to common trait variation between individuals. A high motivation to map such variants is shared worldwide because many heritable traits relate to social and economical preoccupations, such as human health or agronomical and industrial yields. In addition to the molecular knowledge they provide, these variants fuel the development of personalized and predictive medicine as well as the improvement of economically-relevant plants, animal breeds or biotechnology materials. However, this high ambition is accompanied by a major challenge: common traits are under the control of numerous variants that each contribute little to phenotypic variation [1], and this modest contribution of each variant hampers the statistical power to detect them. Power is further limited by the multiplicity of linkage tests when scanning whole genomes. The consequence of this has been debated under the term “missing heritability”: most of the genetic variants of interest remain to be identified. Currently, the only way this issue is handled is by scaling up sample size. For example, human Genome-Wide Association Studies (GWAS) now typically recruit tens of thousands of individuals in a hunt for small-effect variants [2-4]. Practically, however, cohort size cannot be infinitely increased. Studies would therefore greatly benefit from a better detection of small genetic effects, and from a reduction of the number of genomic loci to test.

A particular class of small-effect variants is associated with predisposition (or incomplete penetrance): carriers of a mutation display a phenotype at increased frequency, but not all of them do. From a deterministic point of view, this can be attributed to factors that modify the expressivity of a mutation. Such modifiers can be genetic or environmental factors that interact with the mutation, or other factors such as gender, age or lifestyle. Genetic mapping is improved when these modifiers are accounted for, but this implies that they are known or, at least, anticipated *a priori*. An alternative point of view is to consider the trait as inherently probabilistic. This is likely appropriate for traits corresponding to tissue or cellular failures that depend on various parameters. Although their occurrence in one individual cannot be predicted, risk differences between individuals can be investigated. Under this probabilistic point of view, the statistical properties of cellular traits become interesting: a tissue may break because cells have an increased probability to detach, a tumor may emerge because a cell type has an increased probability of somatic mutations, a chemotherapy may fail if cancer cells have an increased probability to be in a persistent state. In other words, molecular events in one or few cells can have devastating consequences at the multicellular level. Relating single-cell properties to tissular or macroscopic traits can be made by investigations at the molecular level, such as stochastic profiling [5], and at the cellular level, such as recording the response of cell populations to treatment or differentiation signals [6,7].

As discussed previously [8], cellular-scale probabilities are likely related to the genotype and this relation may sometimes underlie genetic predisposition [9]. As a striking example, the genetic factors affecting the mutation rate of somatic divisions are the focus of intense research, because they modify cancer predisposition. These loci have a probabilistic effect on a cellular trait: the amount of *de novo* mutations in the cell’s daughter. Another example is the heterogeneity between isogenic cancer cells which was shown to underlie tumour progression [10,11] and resistance to chemotherapy [6,7,12]. In this context, the fraction of problematic cells may differ between individuals because of their genotypic background. Genetic loci changing the fraction of these cells are likely to modulate disease progression or treatment outcome.

Fortunately, the experimental throughput of single-cell measurements has recently exploded. Technological developments in high-throughput flow cytometry [13], multiplexed mass-cytometry [14], image content analysis [15-17] and droplet-based single-cell transcriptome profiling [18,19] now offer the possibility to estimate empirically the statistical distribution of numerous molecular and cellular single-cell quantitative traits. We therefore propose to scan genomes for variants that modify single-cell traits in a probabilistic manner, which we call single-cell Probabilistic Trait Loci (scPTL). This requires to monitor not only the macroscopic trait of many individuals but also a relevant cellular trait in many cells of these individuals. Including single-cell measurements in genetic association studies would inject high-size samples (thousands of cells per individual) within linkage tests, which could increase detection power while keeping cohort size affordable. After scPTL are found, they can constitute a set of candidate loci to be tested for a possible small effect on the macroscopic trait of interest, thereby avoiding the multiple-testing issue of whole genome association scans.

Methods are needed to detect scPTL. With its fast generation time, high recombination rate and reduced genome size, the unicellular yeast *Saccharomyces cerevisiae* offer a powerful experimental framework for developping such methods. Using this model organism, scPTL were discovered by treating one statistical property of the single-cell trait, such as its variance in the population of cells, as a quantitative trait and by applying Quantitative Trait Locus (QTL) mapping to it [20,21]. However, this approach is limited because it is difficult to anticipate *a priori* which summary statistics must be used.

We present here the development of a genome-scan method that exploits all single-cell values with no prior simplification of the cell population phenotype. Using simulations and existing single-cell data from yeast, we show that it can detect genetic effects that were missed by conventional linkage analysis. When applied to a novel experimental dataset, the method detected a locus of the yeast genome where natural polymorphism modifies cell-to-cell variability of the activation of the GAL regulon. This work shows how single-cell quantitative data can be exploited to detect probabilistic effects of DNA variants. Our approach is conceptually and methodologically novel in quantitative genetics. Although we validated it using a unicellular organism, it opens alternative ways to apprehend the genetic predisposition of multicellular organisms to certain complex traits.

## RESULTS

### Definitions

We specify here the concepts and definitions that are used in the present study. Let *X* be a quantitative trait that can be measured at the level of individual cells. *X* is affected by the genotype of the cells and by their environmental context. However, even for isogenic cells sharing a common, supposedly homogeneous environment, *X* may differ between the cells. To describe the statistical distribution of *X* among cells sharing a common genotype and environment, we define a *single-cell quantitative trait density function f* [8] as the function underlying the probability that a cell expresses *X* at a given level (Figure 1A). Statistically speaking, *f* represents the probability density function of the random variable *X.* In the present study, this function *f(X)* constitutes the ’phenotype’ of the individual from whom the cells are studied. As for any macroscopic phenotype, it can depend on the environmental context of the individual (diet, age, disease…) as well as on its genotype. Single-cell trait density functions also obviously depend on the properties of the cells that are studied, such as their differentiation state, proliferation rate or their capacity to communicate together.

**Figure 1.**
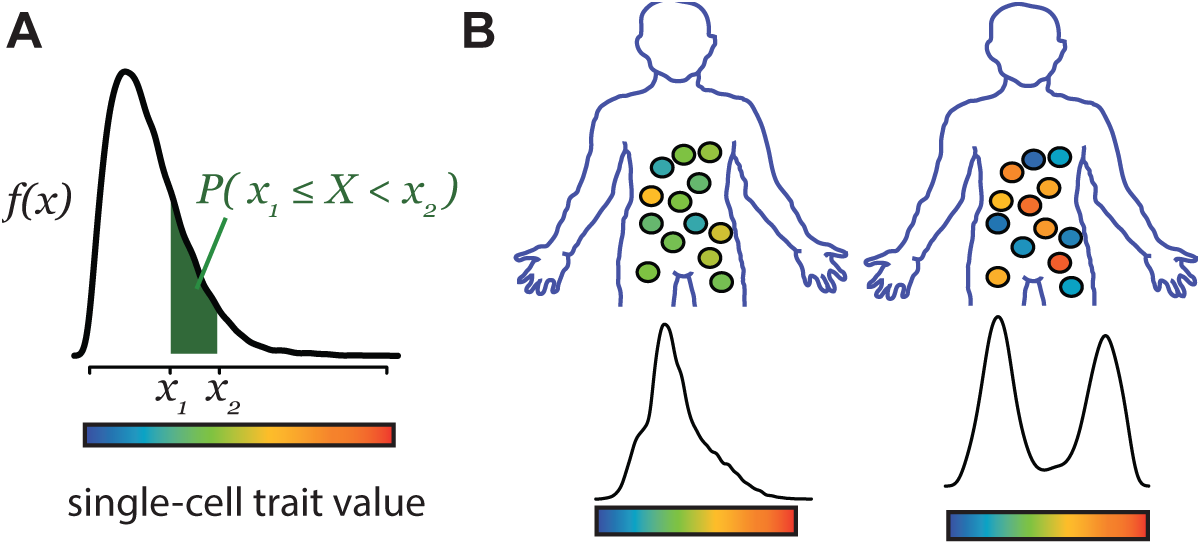
Concept and definitions. **A)** A cellular trait is considered as a random variable *X* with density function *f*. The probability 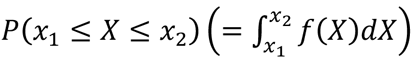 that one cell expresses *X* at a value comprised between *x*_*1*_ and *x*_*2*_ is given by the shaded area. **B**) *f* differs between individuals because of environmental and genetic factors.

We focus here on the effect of the genotype. Conceptually, cells from one individual may follow a density function of *X* that is different from the one followed by cells of another individual, because of genotypic differences between the two individuals (Figure 1B). The important concept is that the genetic difference has probabilistic consequences: it changes the probability that a cell expresses *X* at a given level, but it does not necessarily change *X* in most of the cells. Depending on the nature of trait *X* and how the two functions differ, such a genetic effect can have implications on macroscopic traits and predisposition to disease [8]. The term *single-cell Probabilistic Trait Locus* will refer here to a genetic locus modifying *any* characteristics of *f* (that is, changing allele A in allele B at the locus changes the density function *f* of *X*, i.e. *f*_*B*_ ≠ *f*_*A*_).

A quantitative trait locus (QTL) linked to *X* is a genetic locus that changes the mean or the median of *X* in the cell population. Similarly, a varQTL is a genetic locus changing the variance of *X* and a cvQTL is a genetic locus changing the coefficient of variation (standard deviation divided by the mean, abbreviated CV) of *X* in the cell population. All three types of loci (QTL, varQTL and cvQTL) assume a change in *f* and they are therefore special cases of scPTL. However, not all scPTL are QTL: many properties of *f* may change while preserving its mean, median, variance or CV. The purpose of the present study was to develop an approach that could identify scPTL without knowing *a priori* how it might change *f*.

### Mean and variability of cellular traits have distinct genetic heritabilities

An important question before investing efforts in scPTL mapping is whether genotypes frequently modify *f* without affecting its expected value (the mean of *X*). If this is rare, then QTL mapping will, in most cases, capture the genetic modifiers of *f* and searching for more complex scPTL is only occasionally justified. In contrast, if other-than-mean genotypic changes of *f* are frequent, then scPTL can considerably complement QTL to control single-cell traits. In this case, scPTL mapping becomes important.

In multicellular organisms, cell types and intermediate differentiation states constitute the predominant source of cellular trait variation. Studying their single-cell statistical characteristics requires accounting for the developmental status of the cells. This constitutes a major challenge that can be avoided by studying unicellular organisms. The yeast *S. cerevisiae* provides the opportunity to study individual cells that all belong to a single cell type, in the context of a powerful genetic experimental system. By analysing specific gene expression traits in this organism, we and others identified loci that meet the definition of scPTL but not of QTL [21,22]. This illustrated that, for some traits, scPTL mapping could complement classic quantitative genetics to identify the genetic sources of cellular trait variation.

To estimate if non-QTL scPTL are frequent, we re-analysed an experimental dataset corresponding to the genetic segregation of many single-cell traits in a yeast cross (Figure 2A). After a round of meiosis involving two unrelated natural backgrounds of *S. cerevisiae*, individual segregants had been amplified by mitotic (clonal) divisions and traits of cellular morphology were acquired by semi-automated fluorescent microscopy and image analysis [23]. This way, for each of 59 segregants, 220 single-cell traits were measured in hundreds of isogenic cells, which enabled QTL mapping of these traits. We reasoned that if all scPTL of a trait are also QTL, then a high genetic heritability of any property of *f* should coincide with a high genetic heritability of the expected value of *f*. In particular, the coefficient of variation (CV) of a single-cell trait should then display high heritability only if the mean value of the trait also does. To see if this is the case, we computed for each trait the broad-sense genetic heritabilities of both the mean and CV of the trait. Note that the genetic heritability computed here is not the same as the mitotic heritability of cellular traits transmitted from mother to daughter cells. Here, a value (mean or CV) is computed on a population of cells, and its heritability corresponds to the proportion of its variation that can be attributed to genetic differences between the cell populations (see methods). Strikingly, the two types of heritabilities were not correlated, with many cases of traits showing high heritability of CV and low heritability of their mean value, and *vice versa* (Figure 2B). This indicates that, for many traits, genetic factors exist that modify the trait CV but not the trait mean. This observation is in agreement with the complex CV-vs-mean dependency previously reported in this type of data [24,25]. We therefore sought to develop a method that can detect scPTL that do not necessarily correspond to QTL.

**Figure 2.**
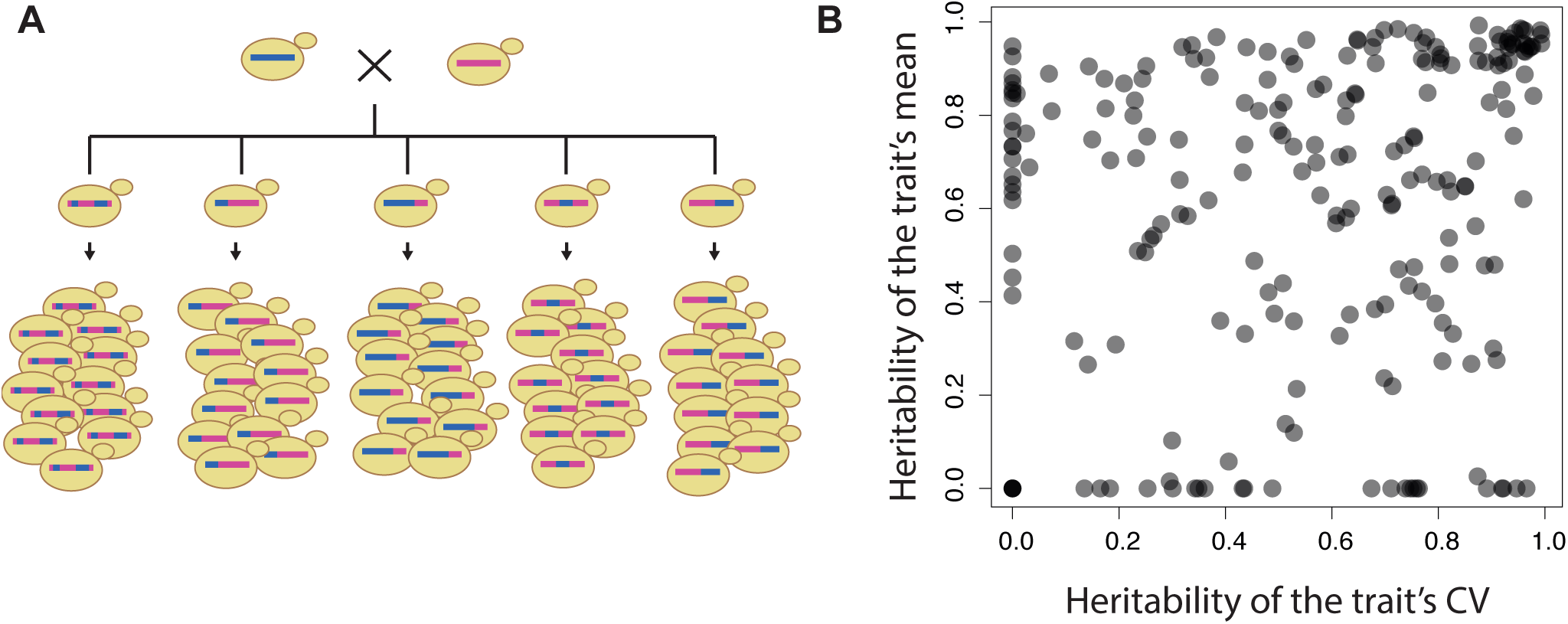
Genetic heritabilities for mean and CV of cellular yeast traits are not correlated. **A)** Scheme of the experimental data used to compute genetic heritabilities. The dataset is from [23]. **B**) Broad-sense genetic heritability of mean (y-axis) and CV (coefficient of variation, x-axis) for 220 traits describing the morphology of individual cells (see methods). Each dot corresponds to one trait. Negative values were set to zero.

### Principle of scPTL mapping

One way to identify scPTL from experimental measures is to compute a summary statistics of the trait distribution, such as one of its moment, and then scan for QTL controlling this quantity. This approach is particularly appropriate when searching for specific genetic effects on *f*, such as a change in the level of cell-to-cell variability, and a few previous studies successfully used it to map varQTL and cvQTL [20,21,23,26,27]. However, it is less adapted when nothing is known on the way *f* may depend on genetic factors.

Scanning for scPTL considers the entire distribution of single-cell trait values as the phenotype of interest and searches the genome for a statistical association with *any* change in the distribution. We assume that for a set of genotypic categories (individuals for multicellular organisms, or populations of cells for unicellular ones), a cellular trait has been quantified in many individual cells of the same type. This way, the observed distribution of the trait constitutes the phenotypic measure of individuals. We also assume that a genetic map is available and the individuals have been genotyped at marker positions on the map. The method we propose is based on three steps. First, a distance is computed for all pairs of individuals in order to quantify how much their phenotype differs. We chose the Kantorovich metric to measure this distance because, unlike the Kullback-Leibler divergence, it satisfies the conditions of non-negativity, symmetry and triangle inequality and, unlike the Hellinger distance, it does not converge to a finite upper limit when the overlap between distributions diminishes [28]. The Kantorovich metric (also called earth-moving distance) can be viewed as the minimum energy required to redistribute one heap of earth (one *f*-function) into another heap (a second *f*-function). It has enabled developments in various fields, ranging from mathematics [29] to economy (the minimal transportation problem) [30,31] to the detection of states from molecular dynamics data [28]. The next two steps are inspired from methods used in ecology, where spatial distinctions between groups are often searched after determining distances between individuals [32,33]. In step 2 of our method, individuals are placed in a vectorial space while preserving as best as possible the distance between them (Figure 3A). This is achieved by multi-dimensional scaling, a dimension-reduction algorithm [34]. The third step is the genetic linkage test itself. At every genetic marker available, a linear discriminant analysis is performed to interrogate if individuals of different genotypic classes occupy distinct sectors of the phenotypic space (Figure 3B-C). Note that if the dimensions have been reduced to a single one, then canonical analysis is not needed: the phenotypic value of each individual has become a scalar and linkage can be performed by standard QTL mapping. Finally, scPTL linkage is scored using the Wilks’ lambda statistics. Statistical inference is made using empirical *p*-values produced by permutations where the identities of individuals are re-sampled. The full procedure is described in details in the methods section.

**Figure 3.**
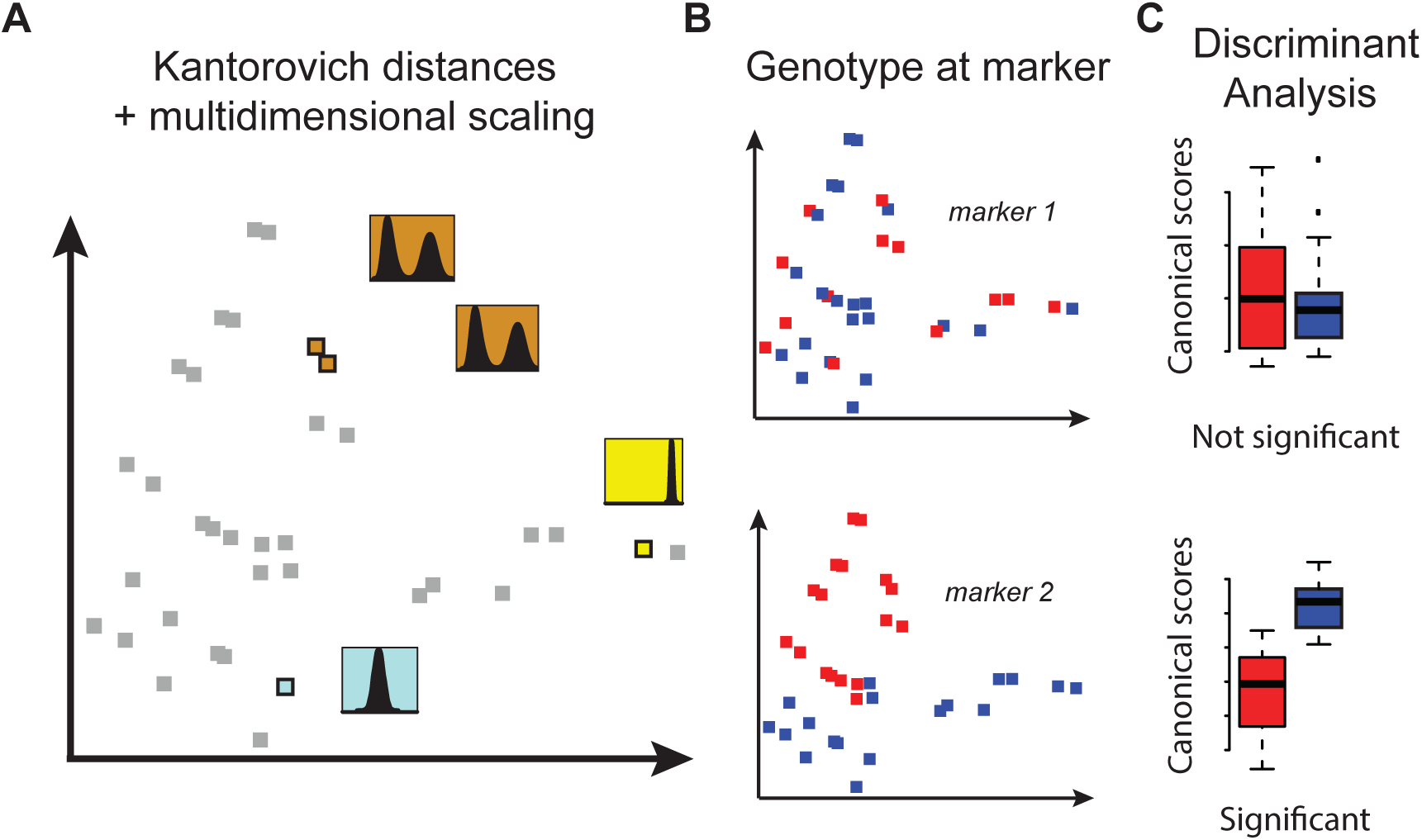
Principle of scPTL mapping. A cohort of multi-cellular individuals (or unicellular clones) with differing genotypes is used. For each individual (or clone), a cellular trait *X* is measured on a population of cells, and the observed distribution of *X* corresponds to the ’phenotype’ of the corresponding individual. **A**) Kantorovich distances are computed for all pairs of individuals. The resulting distance matrix is used to place individuals in a multidimensional space. Proximity of individuals (grey and colored squares) in this space reflects comparable phenotypes (distributions in insets). **B**) Individuals are ’labeled’ (blue *vs.* red) by their genotype at one genetic marker. **C**) A canonical discriminant analysis is performed to test if the genotype at the marker discriminates individuals in the phenotypic space. In the examples displayed, genetic linkage is significant at marker *2* but not at marker 1.

### Detection of simulated scPTL

We first evaluated if our method could detect scPTL from simulated datasets. To do this, we considered a probabilistic single-cell trait governed by a positive feedback of molecular regulations. This is representative of the expression level of a gene with positive autoregulation. As depicted in Figure 4A, the employed model is based on three parameters. For each individual, a set of parameter values was chosen and single-cell values of expression were generated by stochastic simulations. We chose to simulate a scPTL that modified the expected values of the parameters. To do so, we considered a panel of individuals and their genotype at 200 markers evenly spaced every 5cM. Parameter values of each individual were drawn from Gaussians and the mean of these Gaussians depended on the genotype at the central marker. This defined two sets of phenotypes that are depicted by blue and red histograms in Figure 4B. A universal noise term *η* was added to introduce intra-genotype inter-individual variation which, in real datasets, could originate from limited precision of measurements or from non-genetic biological differences between individuals. For each of five increasing values of η, about 130 individuals were simulated.

**Figure 4.**
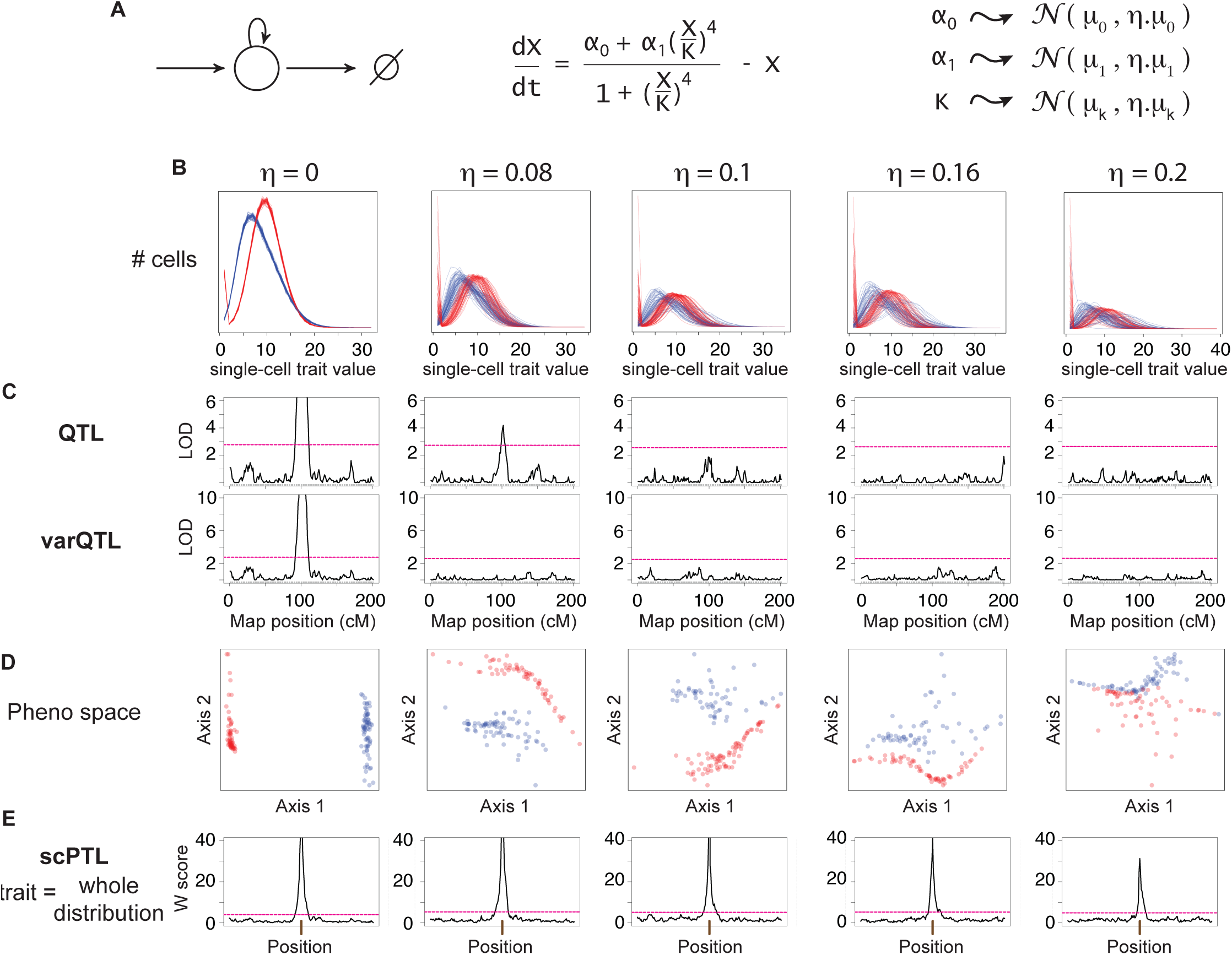
Test on simulations. **A**) A model of gene expression with positive feedback was used to simulate data (see methods). For each of ~130 distinct individuals, parameters *α*_0_, *α*_1_ and *K* were drawn from Gaussian distributions and then used to generate independent values of *X* in 10,000 cells of each individual. Mean values μ_0_, μ_1_ and μ_K_ depended on the genotype of individuals at a locus located in the middle of a 200cM genetic map. Other sources of inter-individual variability were modeled by the extrinsic noise strength η. **B**) Distributions obtained (one per individual) at various values of η. Color: genotype at the locus controlling μ_0_, μ_1_ and μ_K_. **C**) QTL scans. For each individual, the mean (upper panels) or the variance (lower panels) of X were considered as quantitative traits and the map was scanned using interval mapping. Red dashed line: genome-wide significance threshold at 0.05. **D**) Coordinates of individuals (dots) in the phenotypic space obtained after computing Kantorovich distances and applying multi-dimensional scaling. Only the first two dimensions are shown. **E**) scPTL scan. At every marker position, linear discriminant analysis was performed. W score: −log_10_(*Λ*), where *Λ* is the Wilks’ lambda statistics of discrimination (see methods). Red dashed line: empirical genome-wide significance threshold at 0.05 (see methods).

We first scanned the generated dataset by QTL mapping, treating either the mean trait or its variance as the phenotype of interest. This way, the central scPTL locus was detected only when intra-genotype noise was null or very low (Figure 4B). This was anticipated because the mean and variance of the simulated trait values slightly differed between the two sets of individuals. In contrast, our new method allowed to robustly detect the scPTL locus even in the presence of high (up to 20%) intra-genotype noise (Figure 4D-E).

### scPTL mapping of yeast cell morphology

The results described above using a simulated dataset suggest that the method can complement usual QTL mapping strategies. To explore if this was also the case when using real experimental data, we applied scPTL scans to the dataset of Nogami et al. [23] mentioned above (Figure 2A) where 220 single-cell traits were measured in hundreds of cells from segregrants of a yeast cross. We applied three genome x phenome scans, each one at FDR = 10%. Two consisted of QTL interval mapping and were done by considering either the mean cellular trait value of the population of cells or the coefficient of variation of the cellular trait as the population-level quantitative trait to be mapped. The third scan was done using the novel method described here to map scPTL. Significant linkages obtained from this scan are available in Table S1. As shown in Figure 5, the three methods produced complementary results. We detected more linkages with the scPTL method than with the 2 QTL scans combined (71 vs. 61 traits mapped). This illustrates the efficiency of using the full data (whole distribution) of the cell population rather than using a summary statistic (mean or CV). In addition, we expected that a fraction of scPTL would match QTL, because QTL controlling the mean or CV of cellular traits are specific types of scPTL. This was indeed the case, with 67% of scPTL corresponding to loci that were detected by at least one of the two QTL scans. For 11 cellular traits, a locus was found by QTL or cvQTL mapping but it was missed by the scPTL scan. This illustrates that the methods have different power and sensitivity. Importantly, 22 cellular traits were associated to scPTL that were not detected by the QTL search, suggesting that some probabilistic effects may affect poorly the trait’s mean or CV. Altogether, these observations highlight the complementarity of the different approaches and show that scPTL mapping can improve the detection of genetic variants governing the statistical properties of single-cell quantitative traits.

**Figure 5.**
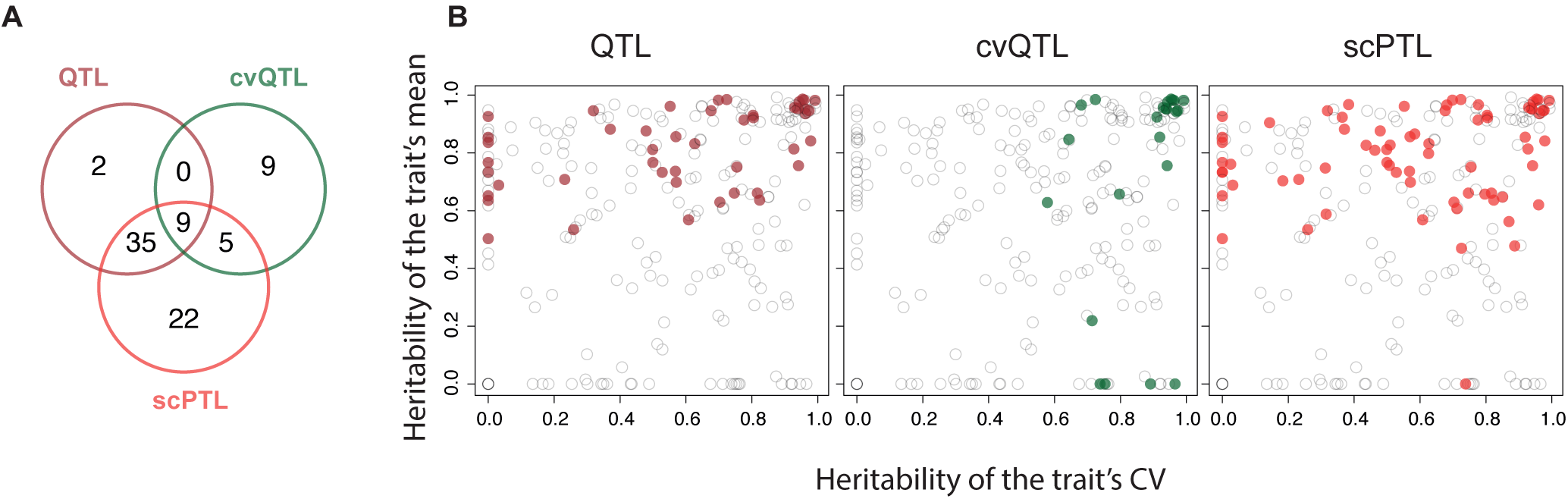
Complementarity of scPTL and QTL mapping using experimental data. The data of Nogami et al. [23] was used to perform genomic scans for QTL, cvQTL and scPTL. For scPTL mapping, we used both the first-axis only and multiple dimensions of the phenotypic space (see methods), and the results were pooled. **A**) Venn diagram showing the number of traits for which a significant locus was found in the genome by each method (each at FDR = 10%). **B**) Same representation as Figure 2B, with colored dots corresponding to the traits mapped by the indicated method.

Examples of scPTL of yeast cellular morphology are shown in Figure 6. One of the cellular traits measured was the distance between the center of the mother cell and the brightest point of DNA staining (Figure 6A). No QTL was found when searching genetic modifiers of the mean or CV of this trait, but a significant scPTL was mapped on chromosome II. When displaying trait distributions, it was apparent that segregants carrying the BY genotype at the locus had reduced cell-cell variability of the trait as compared to segregants having the RM genotype (Figure 6A, right panel). Consistently, a small cvQTL peak was seen on chromosome II, although this peak did not reach genome-wide statistical significance. This trait, which relates to the statistical properties of DNA migration during the early phase of cell division, provided a biological example where scPTL scan identified a genetic modulator of cell-to-cell variability that was missed by the QTL approach.

**Figure 6.**
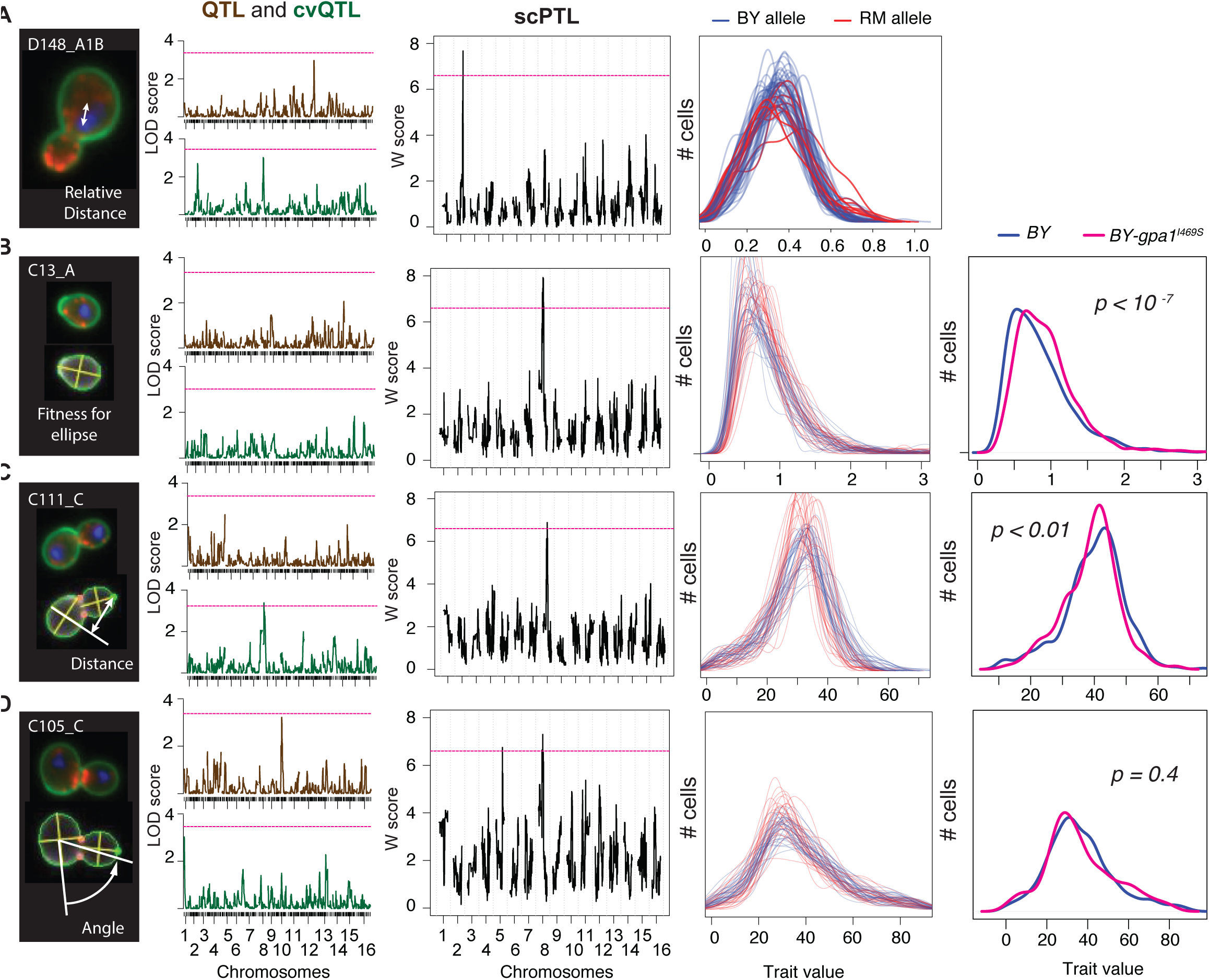
Mapping single-cell probabilistic traits of cellular morphology. For each of 59 recombinant BYxRM yeast strains, four quantitative traits were measured on >600 individual cells [23]. From left to right: description of the trait; results from QTL scans applied to the mean (brown) or coefficient of variation (green) of the trait; results from scPTL scan (pink dashed line: genome-by-phenome significance threshold at FDR = 10%; single-cell trait density functions computed from the data, where each line corresponds to a recombinant strain (color: genotype at scPTL); when relevant, single-cell trait density functions of nearly-isogenic BY strains differing for one non-synonymous SNP in the *GPA1* gene are shown (*p*: statistical significance of the corresponding two-sample Kolmogorov-Smirnov test). **A**) Trait D148_A1B is the distance between the nuclear brightest point and the mother center, relative to the mother size. An scPTL was detected on chromosome 2. **B**) Trait C13_A is the fitness of the cell outline to the best adjusted ellipse. **C**) Trait C111_C is the distance between the bud tip and the extension of the mother short axis. **D**) Trait C105_C corresponds to the position of the budding site. It is the angle between the long axis of the mother cell and the line defined by the mother center and the middle point of neck.

Three other traits were of particular interest because they mapped to a position on chromosome VIII where a functional SNP was previously characterized in this cross. This SNP corresponds to a non-synonymous I->S mutation at position 469 in the GPA1 gene. It causes a residual activation of the pheromone response pathway in the BY strain, and it is associated with an elongated shape of the cells of this strain [23,35]. Here we saw that this locus is an scPTL, but not a QTL, of the degree to which cells are elliptical (Figure 6B). This is consistent with the presence of infrequent ’shmooing’ (elongation that normally takes place in response to pheromone) in the BY strain. Displaying the distributions of this trait in each segregant revealed a remarkable amount of variability between the segregants, and that the BY allele at the locus corresponded to a modest reduction of the trait value as compared to the RM allele (sharper mode at slightly lower value). To see if this was due to the GPA1^I469S^ mutation, we examined the data from a BY strain where this mutation was cured [23]. Remarkably, the single amino-acid substitution caused a mild but statistically significant redistribution of the trait values (Figure 6B). This change was comparable to the difference seen among the segregants, demonstrating the causality of the GPA1^I469S^ SNP. Another trait, corresponding to the distance between the bud tip and the short axis of the mother cell, also mapped to this locus, with the RM allele associated to greater cell-cell variability, and data from the GPA1^I469S^ allele-replaced strain validated this SNP as the causal polymorphism (Figure 6C). These results illustrate that scPTL scans can identify individual SNPs that modify single-cell trait distributions without necessarily affecting the trait mean.

Finally, another trait corresponding to the angle of bud site position mapped to two scPTL loci and no QTL. One of these loci was the GPA1 locus on chromosome VIII. Although the phenotype of bud site selection is not related to ’shmooing’, we examined if the GPA1^I469S^ SNP was involved and found that it was not: the allele-replaced strain did not show a different trait distribution than its control (Figure 6D). Thus, other genetic polymorphisms at the locus should participate to the statistical properties of cellular morphology, by affecting the position of budding sites.

### scPTL scanning detects a new yeast locus that modulates cell-cell variability of the transcriptional response to galactose

We then explored if scPTL scanning could provide new results when applied to a molecular system that had been extensively characterized by classical genetics. The system we chose was the yeast GAL regulon which, in addition to be one of the best described regulatory network, presented several advantages. Natural strains of *S. cerevisiae* are known to display differences in its regulation [36,37] and the transcriptional response of cell populations can be tracked by flow cytometry. This provides data from large numbers of cells and therefore a good statistical power to compare single-cell trait distributions. In addition, acquisitions on many genotypes are possible using 96-well plates. We reasoned that if features of the cell population response segregate in the BY x RM cross (described above for morphology), then scPTL scanning might identify genetic variants having non-deterministic effects on the regulation of GAL genes.

We first compared the dynamics of transcriptional activation of the network in the two strains BY and RM. This was done by integrating a P_Gal1_-GFP reporter system in the genome of the strains, stimulating them by addition of galactose in the medium, and recording the response by flow cytometry. As shown in Figure 7, both strains responded and full activation of the cell population was reached after ~2 hours of induction. Interestingly, remarkable differences were observed between the two strains regarding the distribution of the cellular response. The BY strain showed a gradual increase of expression through time that was relatively homogeneous among the cells (unimodal distribution with relatively low variance), whereas the RM strain showed elevated cell-cell heterogeneity at intermediate activation time points (higher variance, with fraction of non-induced cells). This suggested that genetic polymorphisms between the strains might control the level of heterogeneity of the cellular response at these intermediate time points.

**Figure 7.**
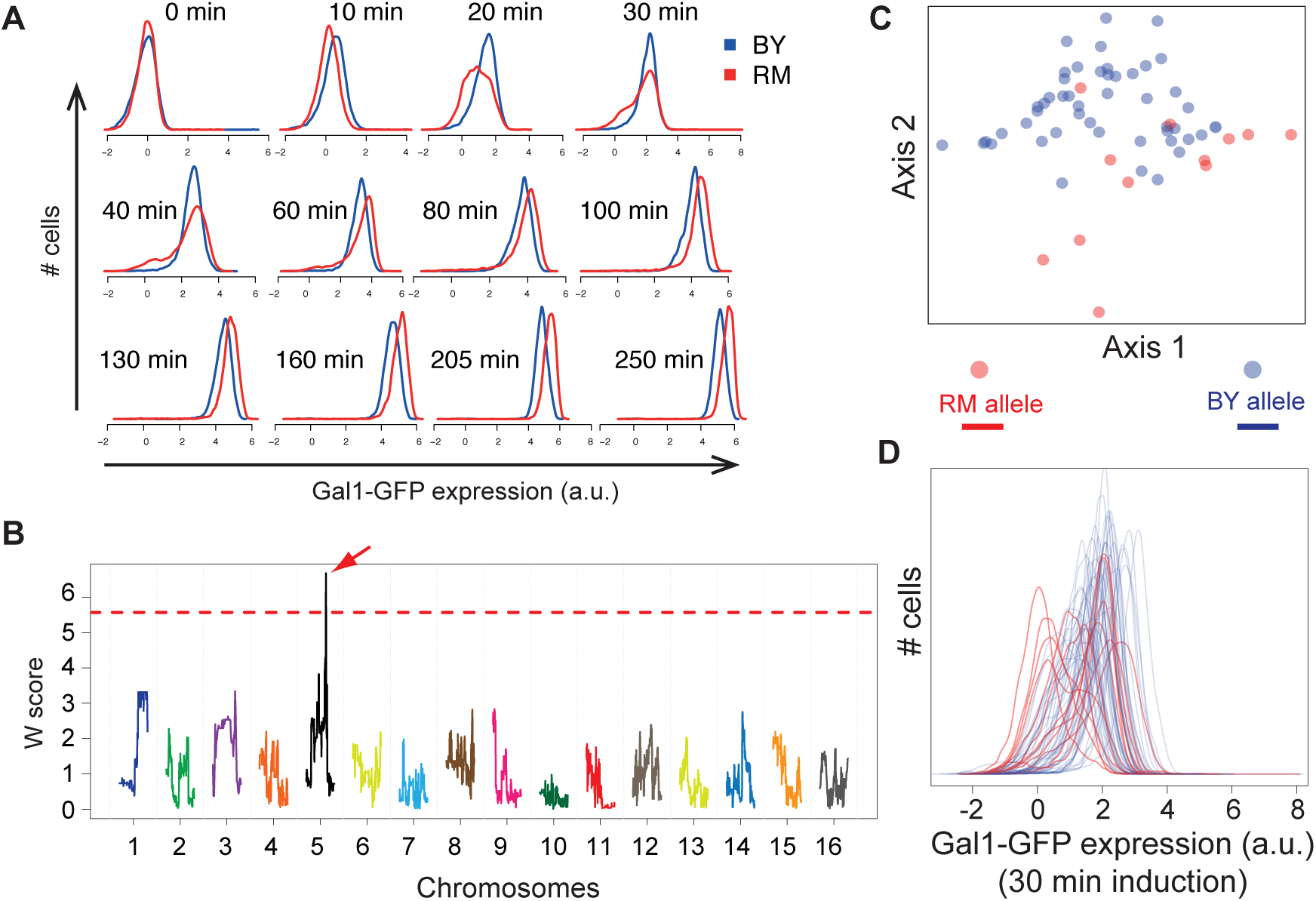
Detection of a scPTL for the cellular response to galactose. **A**) Time-course flow cytometry acquisitions of the response to galactose in strains BY and RM. Cells were cultivated in raffinose 2% and were shifted to a medium containing Raffinose 2% and Galactose 0.5%. After the indicated time, cultures were fixed with paraformaldehyde and analysed by flow cytometry. Histograms correspond to the fluorescent values obtained on cells gated for cell-size (see methods). **B**) Genome scan for scPTL affecting the response after 30 minutes induction. Data similar to panel A was generated for 60 segregants, and the histograms obtained at 30 min post-induction (shown in D), together with the genotypes from [57] were used for scPTL mapping using the multi-dimensions method. The linkage profile (W score) obtained when retaining the first two dimensions is shown, colored by chromosome. Dotted line: significance threshold at genome-wide *p*-value < 0.005. Arrow: significant scPTL on chromosome 5. **C**) Two-dimensional coordinates of the 60 segregants in the phenotypic space (30 min induction time). color: genotype at the scPTL locus. **D**) Phenotypes (histograms of single-cell expression value) of the 60 segregants after 30 min induction, colored by the genotype of the segregants at the scPTL locus. a.u.: arbitrary unit.

We sought to map one or more of these genetic factors. To do so, we acquired the response of 60 meiotic segregants of the BY x RM cross. Using the data collected at each time point, we scanned the genome for scPTL of the reporter gene expression level using the novel genome-scan method described above. The procedure identified a locus on chromosome V position 350,744 that was highly significant (genome-wide *p*-value < 0.001) at 30 minutes post induction, the time at which heterogeneity markedly differed between the BY and RM strains (Figure 7B-C). The locus was also significant at times 20 min (p < 0.005) and 40 min (p < 0.005) post induction.

Visualizing the distributions of single-cell expression levels at 30 minutes revealed that the RM and BY genotypes at this locus corresponded to high and low cell-cell heterogeneity, respectively (Figure 7D). Thus, this locus explains, at least in part, the different levels of heterogeneity observed between the parental strains. It should therefore also be detected as a varQTL or cvQTL. This was indeed the case: the LOD score linking the locus to the variance of expression was 4.5 and reached statistical significance (*P* = 0.005). Importantly, the scPTL was not a QTL: the locus genotype did not correlate with the mean level of expression of the population of cells (LOD score < 2.8).

When surveying the genomic annotations of the locus [38], we realized that it contained no obvious candidate gene that would explain an effect on the heterogeneity of the response (such as genes known to participate to the transcriptional response). One potentially causal gene was DOT6, which encodes a poorly characterized transcription factor that was shown to shuttle periodically between the cytoplasm and nucleus of the cells in standard growth conditions [39]. Given that i) the shuttling frequency of such factors can sometimes drive the response to environmental changes and ii) numerous non-synonymous BY/RM genetic polymorphisms were present in the gene, we constructed an allele-replacement strain for DOT6 and tested if the gene was responsible for the scPTL linkage. This was not the case. Strains BY and BY-*DOT6*^*RM*^ (isogenic to BY except for the DOT6 gene which was replaced by the RM allele) displayed very similar transcriptional responses at intermediate times of induction (Supplementary Figure 1). Fine-mapping of the locus and a systematic gene-by-gene analysis are now needed to precisely identify the polymorphisms involved. By highlighting a novel genetic locus modulating cell-cell variability of the transcriptional response to galactose, our results show that scPTL scanning can provide new knowledge on the fine structure of a well-studied system.

## DISCUSSION

We have developed a novel method to scan genomes for genetic variants affecting the probabilistic properties of single-cell traits. We validated the method using data from colonies of a unicellular organism, which constitutes a first step before transferring the method to multicellular organisms. Our approach extends the usual genetic analysis of quantitative traits both conceptually and methodologically: by incorporating large samples of phenotypic values at the cellular scale, variants that have probabilistic effects can be detected and their possible contribution to trait heritability at the macroscopic (multicellular) scale can be investigated. In this section, we first compare our method to existing ones, we then consider how scPTL detection can be further improved or extended, and we finally discuss the biological relevance of scPTL mapping to cancer predisposition and other complex traits.

### Methods for scPTL mapping

Our new method based on the Kantorovich distance is not the only one by which scPTL can be identified. Applying classical QTL mapping to summary statistics of the cellular traits can also be efficient. We emphasize that the two approaches are complementary. For example, our method missed to detect linkage for 9 yeast morphological traits for which cvQTL scans were successful, but it detected several significant scPTL that were missed by the QTL-based approach (Figures 5 and 6). Second, we observed that scPTL detection was often efficient when the mean value of cellular trait differed among genotypic categories. As shown on Figure 6B, traits successfully mapped tended to display high heritability of the mean. Thus, after a scPTL is detected, it is necessary to examine the effect on the trait distributions and to determine if it is a QTL or not. Third, alternative ways of mapping scPTL are open and may prove more appropriate in some contexts. For example, if a cellular trait becomes preoccupying when it exceeds a certain threshold value, then the fraction of cells above this threshold can be used as a macro-trait to be mapped by QTL analysis. This way, the focus is made on the relevant aspect of the cellular trait, avoiding variation in other parts of the distribution. We therefore recommend conducting Kantorovich-based scPTL mapping in addition to classical methods and not as a replacement strategy.

### Future improvements in scPTL mapping

While the principle of genotype-phenotype genetic linkage dates back to several decades ago, the statistical methods that test for linkage are still being improved, especially regarding multi-loci interactions or population structure corrections [40,41]. The present study provides a priming of a generic scPTL mapping approach (exploiting thousands of single-cell trait values) and demonstrates its feasibility and potential (new loci were detected). Since it is new, we anticipate that it will also evolve in the future. It is currently based on three steps: (i) compute pairwise distances between individuals by using the Kantorovich metric, (ii) use the resulting distance matrix to construct a relevant phenotypic space and (iii) test for genetic linkage by LDA. A number of methodological considerations can be made in anticipation to future developments and applications.

A phenotypic space can be constructed by alternative ways that do not require the Kantorovich metric. For example, we considered representing individuals in a “space of moments”, where the coordinate of every individual on the i-th axis was the i-th moment of the cellular trait distribution associated to this individual. We applied this to the yeast morphological data and we searched for genetic linkages by linear discriminant analysis as described above. This approach detected many significant scPTL but we encountered a difficulty that was avoided by our Kantorovich-metric based method. When searching for significant linear discrimination, the dimensionality of the phenotypic space is important. At high dimensionality, discriminant axes are more likely to be found. This improves detection in the actual data but at the expense of increasing the degrees of freedom and therefore the false positive hits estimated from the permuted data. In a "space of moments", the properties of the single-cell trait distributions are very important because they define which axis (moments) are relevant to separate individuals. Keeping the 4-th axis may be crucial even if all individuals have very similar first, second or third moments. Choosing the appropriate dimension for LDA is then arbitrary and it becomes difficult to keep a good detection power while still controlling the FDR. This issue is avoided in the case of Kantorovich distances because multi-dimensional scaling can be applied without normalization and the axes of the phenotypic space are ranked by descending order of their contribution to the inter-individual differences. The 4-th axis, for example, contributes less than the first three axes to the separation of individuals in the space. If keeping the 4-th dimension prior to LDA is beneficial for linkage, then keeping the first three axes is also highly relevant, and this is true regardless of the properties of the single-cell trait distributions. We found this very useful: our algorithm adds dimensions one by one and evaluates the benefit of each increase (see methods).

There are at least three lines along which our method may be further improved. First, LDA is only appropriate if genotypic categories can be distinguished along linear axis. If individuals in the phenotypic space are separated in non-linear patterns, other methods such as those based on kernel functions [42] may be more appropriate. Second, it is currently not clear how confidence intervals of scPTL can be computed. Filling this caveat would help practical studies that need to prioritize the efforts needed to identify the causal DNA polymorphism. And third, single-cell data acquisitions often generate multiple trait values for each individual cells. This is the case for morphological profiling as in the dataset we used here, but also for gene expression [43] or parameters describing the micro-environment of the cells [44]. It would therefore be interesting to search for scPTL affecting multiple cellular traits simultaneously instead of treating cellular traits one by one. A multidimensional analysis could be performed in order to extract a set of informative meta-traits, such as principal components or representative medoids and scPTL of these meta-traits could be searched using our method. This dimension-reduction approach would benefit from the redundant information available from correlated traits (e.g. the perimeter of a cell and its area are two measurements of its size), but the biological interpretation of a probabilistic effect on a meta-trait may not be straightforward. Alternatively, one might want to identify scPTL affecting the joint probability distribution of multiple cellular traits. In this case, a natural extension of our method would be to compute Kantorovich distances between multivariate distributions. However, the Kantorovich metric cannot be easily computed for more than two marginals (i.e. cellular traits in our case). In fact, its existence as a unique solution to the multi-dimensional transportation problem was itself a subject of research [45]. A possible alternative could be to compute a Euclidean distance in the "space of moments" mentioned above and then apply multi-dimensional scaling.

### scPTL and the genetic predisposition to disease

Beyond the methodological aspects discussed above, it is important to distinguish the situations where scPTL mapping is biologically relevant from those where it is not. The determinants of human height, for example, act via countless cells, of multiple types, and over a very long period of time (~ 16 years). In such cases, the macroscopic trait results from multiple effects that are cumulated and considering the probabilistic individual contribution of specific cells is inappropriate. In contrast, a number of macroscopic traits can be affected by particular events happening in rare cells or at a very precise time (see below). In these cases, studying the probabilities of a biological outcome in the relevant cells or of a molecular event within the critical time interval can provide invaluable information on the emergence of the macroscopic phenotype, and scPTL mapping then becomes relevant.

A striking example of such traits is cancer. Genetic predisposition is conferred by variants affecting somatic mutation rates and these loci are special cases of scPTL: the cellular trait they modify is the amount of *de novo* mutations in the cell’s daughter. These variants have classically been identified by genetic linkage of the macroscopic trait (disease frequency in families and cohorts), and their role on the maintenance of DNA integrity was deduced afterwards by molecular characterizations. For a review on the genetics of cancer syndrome predisposition, see [46,47]. New genetic factors with milder effects on somatic mutation rates may be identified directly. If a reporter system of *de novo* mutations can be introduced in a relevant and large population of cells, then the high number of cell measurements may allow to detect loci that modify even slightly the mutation rate. Alternatively, if the functional properties (expression level, phosphorylation status, subcellular localization) of a critical tumor-suppressor gene can be monitored in numerous individual cells, then scPTL mapping, as presented here, may help identify genetic factors that modulate the activity of this gene in a probabilistic manner. Once identified, the association of these loci with the macroscopic phenotype can then be tested without the statistical challenges of whole-genome scans.

In addition, scPTL mapping is also relevant to the non-genetic heterogeneity of cancer cells which was shown to be associated with tumour progression [10,11] and treatment efficiency [6,7,12]. Genetic loci changing the fraction of problematic cells are likely modulators of the prognosis. In principle, the scPTL mapping method presented here has the potential to identify such loci. In practice, however, the key to success will likely reside in the design of experimental acquisitions: what single-cell trait should be measured? Can it be measured in a sufficiently large number of cells? Choice of the trait can be driven by molecular or cellular knowledge on the physiological process involved. For example, the distribution of the biomarker JARID1B (a histone demethylase) in populations of melanoma cells is indicative of an intra-clonal heterogeneity that is important for tumour progression [10], biomarkers CD24 and CD133 can distinguish rare cells that persist anti-cancer drug treatments [7] and multiplexed markers of signalling response can reveal patterns of population heterogeneity that predict drug sensitivity [48]. When relevant markers are not known, a possibility is to screen for them using stochastic profiling [5]. This method interrogates the transcriptomic variability between pools of few cells in order to identify transcripts displaying elevated cell-to-cell variability in specific biological contexts. It allowed the discovery of two molecular states of extracellular matrix-attached cells that can be distinguished by the *jun D proto-oncogene* and markers of TGF-β signalling [11]. Such markers of isogenic cellular subtypes may allow the development of scPTL mapping in humans.

An important statistical requirement to identify scPTL is the abundance of cells on which the probabilistic trait is quantified. For human studies, peripheral blood offers access to many cells but, unfortunately, many internal organs do not. This requirement also implies using technologies where the throughput of quantitative acquisitions is high. This is the case for flow-cytometry and, although at higher costs, for high-content image analysis [15,16] and digital microfluidics [18,19]. For these practical reasons, it is possible that mouse immunological studies will help making progress in mammalian scPTL mapping. For example, the work initiated by Prince et al. [49] describing pre-and post-infection flow-cytometry profiles of F2 offsprings from different mouse strains may provide an interesting pilot framework.

The interest of scPTL mapping is not restricted to cancer biology. Developmental processes and cellular differentiation are also vulnerable to mis-regulations happening in few cells or during short time intervals. Their macroscopic outcome can therefore be affected by probabilistic events at the cellular scale. For example, stochastic variation in the expression of the stem cell marker Sca-1 is associated with different cellular fates in mouse hematopoietic lineages [50], suggesting that genetic factors changing this stochastic variation may impact blood composition. Similarly, embryonic stem cells co-exist in at least two distinct molecular states that are sensitive to epigenetic and reprogramming factors [51]. Genetic variants modulating these factors may change the statistical partitioning of these states. Two observations made on flies remarkably support the existence of natural genetic factors altering developmental processes in a probabilistic manner. The first one is the fact that high levels of fluctuating asymmetry can be fixed in a wild population of *D. melanogaster* under artificial selection [52]. The second one comes from a comparative study of Drosophila species [53]. Embryos of *D. santomea* and *D. yakuba* display high inter-individual variability of expression of the signal transducer pMad at the onset of gastrulation, as compared to *D. melanogaster* embryos. This increased variability was attributed to a reduced activity of the homeobox gene *zerknüllt* thirty minutes before this stage. Very interestingly, it is accompanied by phenotypic variability (inter-individual variance of the number of amnioserosa cells) in *D. santomea* but not in *D. yakuba*. These and other examples [54] illustrate how developmental variability and phenotypic noise can evolve in natural populations. Applying scPTL mapping may allow to dissect the genetic factors responsible for this evolution.

Finally, we can anticipate that gene-gene and gene-environmment interactions also shape the probability density function of cellular traits. Our results on the activation of the yeast galactose network remarkably illustrate this: the effect of the scPTL on chromosome V is apparent only transiently, and in response to a change of environmental conditions. It is tempting to extrapolate that signalling pathways in plants and animals may be affected by scPTL that act at various times and steps along molecular cascades.

In conclusion, our study provides a novel method that can detect genetic loci with probabilistic effects on single-cell phenotypes, with no prior assumption on their mode of action. By exploiting the power of single-cell technologies, this approach has the potential to detect small-effect genetic variants that may underlie incomplete trait penetrance at the multicellular scale.

## METHODS

### Stochastic modeling of a positively auto-regulated gene

Single-cell gene activity was modeled by a stochastic variable *X* that represented the number of proteins in one cell at a given time. Under the model, the dynamics of *X* is controlled by two processes: (1) protein production with rate α and (2) protein degradation with rate *β*. We assume that the gene is positively auto-regulated by a 4-mer complex, meaning that *α* is an increasing function of *X* with a typical Hill-like shape

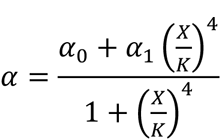

with *α*_*0*_ the leaky production rate in absence of *X* 4-mers at the promoter, *α*_*1*_ the production rate in presence of 4-mers, and *K* the dissociation constant of the 4-mer. We set β=1, which corresponds to scaling time units. The dynamics of the mean value of *X* in a population of isogenic cells follows the equation shown in Figure 4A. To obtain the probability distribution of *X*, we performed exact stochastic simulations of the chemical system defined by the two reactions rate *α* and *β*, using the Stochastic Simulation Algorithm [55].

To generate two groups of individuals, we assumed that the set of parameters (*α*_*0*_, *α*_*1*_, *K)* was controlled by one locus that could exist in two alleles (A and B) with mean values (*μ*_*0*_^*A/B*^, *μ*_*1*_^*A/B*^, *μ*_*K*_^*A/B*^) and, for simplicity, that the individuals were haploids. To account for sources of inter-individual variability within genotypic groups, the values of the parameters for one individual were drawn from normal distributions of mean values *μ*_*0*_^*A/B*^, *μ*_*1*_^*A/B*^ and *μ*_*K*_^*A/B*^ and of standard deviations *ημ*_*0*_^*A/B*^, *ημ*_*1*_^*A/B*^ and *ημ*_*K*_^*A/B*^ where *η* represented the strength of inter-individual variability. *η* was assumed to be the same for A and B alleles.

### Genetic heritabilities of cellular traits’ mean and CV

All statistical analysis were done using R (version 3.1.2) [56]. The data from Nogami et al. [23] consisted of 220 traits, acquired on >200 cells per sample. Note that most traits are related to one of three division stages. Each trait was therefore measured on a subset of cells of the sample (less than 200). There were nine samples of the BY strain, nine of the RM strain, and three of each of 59 segregants of the BY x RM cross. To increase the number of cells per sample (which was helpful for scPTL mapping done afterwards), we pooled triplicates together. The data then corresponded to 220 traits, measured on >600 cells per sample, with 3 BY samples, 3 RM samples, and 1 sample per segregant. For each trait, we computed the genetic heritabilities of the mean and CV as follows. The mean and CV of the cellular trait in each sample were computed, leading to two scalar values per sample that we call macro-traits hereafter (to distinguish them from the single-cell values). The broad-sense genetic heritability of each macro-trait was *H*^*2*^ = (*var*_*T*_ - *var*_*E*_) / *var*_*T*_, where the total variance *var*_*T*_ was the variance of the 59 macro-trait values of the segregants and where the environmental variance *var*_*E*_ was estimated as (*var*_*BY*_ + *var*_*RM*_) / *2*, with *var*_*BY*_ (resp. *var*_*RM*_) being the variance of the three macro-trait values of the BY (resp. RM) strain.

### scPTL mapping of yeast morphological traits

Following [28], we computed the Kantorovich distance between two distributions *f*_*1*_ and *f*_*2*_ as the area under the absolute value of the cumulative sum of the difference between the two distributions:

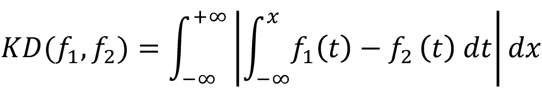

Multi-dimensional scaling of the resulting distance matrix was then performed using the R function `cmdscale()` from the stats package. The number of dimensions retained (*ndim*) was the number of eigenvalues exceeding the expected value under the hypothesis of no structure in the data (i.e. mean of all eigenvalues, Kaiser criterion). We computed the heritability of each yeast morphological trait in this multidimensional space. This was done as above for one dimension, by computing the total variance of the data, and estimating the environmental variance from the replicated experiments made on the parental strains. For 147 traits, heritability was greater than 0.5 and scPTL were searched. Details on how these steps were implemented in R are available in Supplementary Methods.

The yeast genotypes we used were from Smith and Kruglyak [57]. We scanned the genetic map with two methods. First, we considered the coordinates of each segregant on the first axis of the multi-dimensional scaling, and we considered this coordinate as a quantitative trait that we used for interval mapping using R/qtl [58]. Secondly, we applied a linear discrimination analysis (LDA) on the phenotypes data, using the genotype at every marker as the discriminating factor. An important issue in this step is the mutidimensionality of the data: axis 2, 3 and more may contain useful information to discriminate genotypic groups, but if too many dimensions are retained, a highly-discriminant axis may be found by chance only. To deal with this issue, we evaluated the output of LDA at all dimensions *d* ranging from 2 to *ndim*. For each value of *d*, we applied LDA at every marker position and we recorded the Wilks’ lambda statistics:

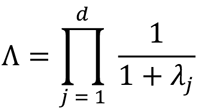

where λ_j_ was the *j*-th eigenvalue of the discriminant analysis. Low values of this statistics allow to reject the null hypothesis of no discrimination by the factor of interest [59] which, in our case, is the genotype. We defined a linkage score (W score) as:

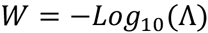

We then quantified how much the best marker position was distinguished from the rest of the genome by computing a Z-score:

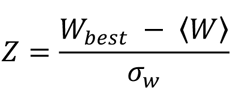

where *W*_*best*_, <*W*> and *σ*_*W*_ were the highest, the mean and the standard deviation of all W scores found on the genome, respectively. Finally, we chose the dimension that maximized this Z-score (i.e. dimension where the linkage peak had highest contrast). Very importantly, the same degrees of freedom (exploration of the results at various dimensionalities) were allowed when applying the permutation test of significance (see below).

### Permutation tests for statistical significance

We first explain the case where a single trait is studied. When the trait was mapped using R/qtl, significance was assessed by the permutation test implemented in function scanone() of the package [58]. For scPTL, we implemented our own permutation test as follows. The significance of an scPTL is the type one error when rejecting the following null hypothesis: "there is no marker at which the genotype of individuals discriminates their location in the phenotypic space", where one ’individual’ refers to one population of isogenic cells, and where the ’phenotypic space’ is the multi-dimensional space built above by computing Kantorovich distances and applying multi-dimensional scaling. The relevant permutation is therefore to randomly re-assign the phenotypic positions to the individuals before scanning genetic markers for discrimination. We did this 1,000 times. Each time, LDA was applied at dimensions 2 to *ndim*, the dimension showing the best contrast (high Z score) was retained, and the highest W score obtained at this dimension was recorded. The empirical threshold corresponding to genome-wide error rates of 0.1%, 1% and 5% were the 99.9^th^, 99^th^ and 95^th^ percentiles of the 1,000 values produced by the permutations, respectively.

We now explain the case of the morphological study, where multiple traits (220) were considered. Four different methods were used. For three of them, single-cell trait values were resumed to a scalar macro-trait and QTL was searched. The three methods differed by the choice of this macro-trait, which was either the mean or the coefficient of variation of single-cell traits, or the coordinate of individuals on the first axis of the phenotypic space. For each of the three methods, morphological traits with less than 50% genetic heritability (see above) were not considered further, and QTL was searched for the remaining *N*_*traits*_ traits only. For each of these traits, LOD scores were computed on the genome by interval mapping using the macro-trait value as the quantitative phenotype of interest. Significance was assessed by random re-assignment of the macro-trait values to the individuals (yeast segregants). We did 1,000 such permutations. For each one, the genome was scanned as above and the highest LOD score on the genome was retained. This generated a 1,000 x *N*_*traits*_ matrix *M*_*perm*_ of the hits expected by chance. At a LOD threshold *T*, the FDR was computed as:

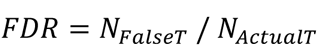

where *N*_*ActualT*_ was the number of linkages obtained from the actual dataset at LOD > *T*, and *N*_*FalseT*_ was the expected number of false positives at LOD > *T*, which was estimated by the fraction of elements of *M*_*perm*_ exceeding *T*.

The fourth method considered all coordinates of the individuals in the phenotypic space. At this step, for each morphological trait, a phenotypic space of *ndim* dimensions had been built as explained above by computing Kantorovich distances and applying multidimensional scaling. Let P1, P2 and PS be the phenotypic matrices of parent 1 (strain BY), parent 2 (strain RM) and segregants, respectively, with rows being the samples (replicates for P1 and P2, and segregants for PS) and columns being the *ndim* coordinates of each sample in the phenotypic space. These matrices had dimensions 3 x *ndim* for P1 and P2 and 59 x *ndim* for PS. Genetic heritability was computed as *H*^*2*^ = (*var*_*T*_ - *var*_*E*_) / *var*_*T*_, where the total variance *var*_*T*_ was the variance of the samples in PS, and where the environmental variance *var*_*E*_ was estimated as (*var*_*P1*_ + *var*_*P2*_) / *2*, with *var*_*P1*_ (resp. *var*_*P2*_) being the variance of the samples in P1 (resp. P2). Morphological traits showing *H*^*2*^ < 0.5 were discarded, and scPTL mapping was applied to the remaining *N*_*traits*_ traits as described above (choice of dimensionality with highest contrast and recording of the best W score obtained on the genome at this dimensionality). Significance of W scores was assessed as described above for the LOD scores, by performing 1,000 permutations and determining the FDR associated to various thresholds of W scores.

### Yeast strains and plasmids

The yeast strains and oligonucleotides used in this study are listed in Table S2.

To construct the Gal-GFP reporter, we first removed the MET17 promoter of plasmid pGY8 [20] by digestion with restriction enzymes BspEI and SpeI followed by Klenow fill-in and religation. This generated plasmid pGY10. The GAL1 promoter fragment was digested (BglII-BamHI) from pFA6a-His3MX6-PGAL1 [60] and cloned in the BamHI site of pGY10. A small artificial open reading frame upstream GFP was then removed by digestion with EcoRV and BamHI, Klenow fill-in end blunting and religation. This generated plasmid pGY37, carrying a *P*_*GAL1*_-*yEGFP-NatMX* cassette that could be integrated at the *HIS3* genomic locus.

Plasmid pGY37 was linearized at NheI and integrated at the *HIS3* locus of strain BY4716 (isogenic to S288c), YEF1946 (a non-clumpy derivative of RM11-1a) and in 61 F1 non-clumpy segregants from BY471xRM11-1a described in [23] to generate strains GY221, GY225, and the S288c x RM11-1a *HIS3:P*_*GAL1*_-*yEGFP-NatMX:HIS3* set, respectively.

In parallel, we also constructed a GAL1-GFP_PEST_ reporter coding for a destabilized fluorescent protein [61]. We derived it from pGY334, where GFP_PEST_ was under the control of the PGK promoter. pGY334 was constructed in several steps. The PGK promoter was PCR-amplified from pJL49 (gift from Jean-Luc Parrou) using primers 1A23 and 1A24, digested by BamHI and cloned into the BamHI site of pGY10. The resulting plasmid was digested with EcoRV and XbaI, subjected to Klenow fill-in end blunting and religated, generating plasmid pGY13 carrying a *HIS3:P*_*PGK*_-*yEGFP-NatMX:HIS3* cassette. The lox-CEN/ARS-lox sequence from pALREP [21] was amplified by PCR using primers 1I27 and 1I28 and cloned by homologous recombination into pGY13, generating plasmid pGY252. The GFP_PEST_ sequence was PCR-amplified from pSVA18 [61] using primers 1I92 and 1I93 and cloned *in vivo* into pGY252 (digested by MfeI and DraIII), leading to pGY334. The GAL1 promoter fragment was amplified by PCR from pGY37 using primers 1J33 and 1I42 and cloned into plasmid pGY334 by recombination at homologous sequences flanking the BamHI site of the plasmid. The CEN/ARS cassette of the resulting plasmid was excised by transient expression of the Cre recombinase in bacteria [21], generating the final integrative plasmid pGY338 carrying the *HIS3:P*_*GAL1*_-*GFP*_*PEST*_-*NatMX:HIS3* cassette.

pGY338 was linearized by NheI and integrated at the *HIS3* locus of BY4724 (isogenic to S288c) and GY1561 to create GY1566 and GY1567 strains, respectively. Strain GY1561 is a non-clumpy derivative of RM11-1a where the KanMX4 cassette was removed. It was obtained by first transforming RM11-1a with an amplicon from plasmid pUG73 [62] obtained with primers 1E75 and 1E76 and selecting a G418-sensitive and LEU+ transformant (GY739) which was then transformed with pSH47 [63] for expression of the CRE recombinase. After an episode of galactose induction, a LEU-derivative was chosen and cultured in non-selective medium (URA+) for loss of pSH47, leading to strain GY744, which was then crossed with GY689 [64] to generate GY1561.

### Galactose response measurements

Liquid cultures in synthetic medium with 2% raffinose were inoculated with a single colony and incubated overnight, then diluted to OD600 = 0.1 (synthetic medium, 2% raffinose) and grown for 3 to 6 hours. Cells were then resuspended in synthetic medium with 2% raffinose and 0.5% galactose and grown for the desired time (0, 10, 20, 30, 40, 60, 80, 100, 130, 160, 205 and 250 minutes). Cells were then washed with PBS1X, incubated for 8 min in 2% paraformaldehyde (PFA) at room temperature, followed by 12 min of incubation in PBS supplemented with Glycine 0.1M at room temperature and finally resuspended in PBS. They were then analyzed on a FACSCalibur (BD Biosciences) flow cytometer to record 10,000 cells per sample.

Flow cytometry data was analysed using the *flowCore* package from Bioconductor [65]. Cells of homogeneous size were dynamically gated as follows: (i) removal of events with saturated signals (FSC, SSC or FL1 ≥ 1023 or ≤ 0), (ii) correction by subtracting the mean(FL1) at t=0 to each FL1 values, (iii) computation of a density kernel of FSC,SSC values to define a perimeter of peak density containing 60% of events, (iv) cell gating using this perimeter and (v) removal of samples containing less than 3,000 cells at the end of the procedure. The GFP expression values were the FL1 signal of the retained cells.

### Dot6 allele replacement

The *DOT6*^*RM*^ allele was amplified by PCR from genomic DNA of the RM strain using primers 1K87 and 1K88. It was then cloned into plasmid pALREP [21] by homologous recombination at sequences flanking the HpaI site of the plasmid. The CEN/ARS cassette of the resulting plasmid was excised by transient expression of the Cre recombinase in bacteria, as previously described [21], generating plasmid pGY389, which was linearized at EcoRI (a unique site within the DOT6 gene) and integrated in strain GY1566 (isogenic to BY, and carrying the *HIS3:P*_*GAL1*_-*GFP*_*PEST*_: *HIS3* cassette). The pop-in pop-out strategy was applied as previously described [21] and four independent transformants were selected (GY1604, GY1605, GY1606 and GY1607) where PCR and sequencing validated the replacement of the DOT6 allele.

### Data Availability

The yeast morphological data corresponds to the experiments described in [23]. For the present study, raw images were re-analyzed using CalMorph 1.0. The single-cell values and genotypes used are available in the supplementary material of this article.

The flow cytometry data corresponding to yeast galactose response is made available from http://flowrepository.org under accession number FR-FCM-ZZPA.

## ACKNOWLEDGEMENTS

We thank Tamiki Komatsuzaki for mentioning the potential usefulness of the Kantorovich metric, Fabien Crauste and Emmanuel Grenier for helpful discussions, Satoru Nogami for re-analysis of CalMorph images, Fabien Duveau, Olivier Gandrillon, Marie Sémon and Gérard Triqueneaux for critical reading of the manuscript, Sandrine Mouradian and SFR Biosciences Gerland-Lyon Sud (UMS344/US8) for access to flow cytometers and technical assistance, the Pôle Scientifique de Modélisation Numérique (Lyon, France) and the Grid’5000 testbed (www.grid5000.fr) for computer resource, and developers of R, bioconductor and Ubuntu for their software.

## FUNDING STATEMENT (TO BE PASTED IN ONLINE FORM)

This work was supported by the European Research Council under the European Union’s Seventh Framework Programme FP7/2007-2013 Grant Agreement n°281359. D.J. was supported by the Institut Rhône-Alpin des Systèmes Complexes and program AGIR of University Grenoble Alpes.

## AUTHORS CONTRIBUTIONS

conceived and designed the experiments: GY

performed the experiments: MR, HDB

analyzed the data: FC, MR, DJ

contributed reagents/materials/analysis tools: FC, DJ, YO

contributed to the writing of the manuscript: MR, GY

other:

Developed the scPTL mapping method: FC and GY. Developed the analysis code: FC. Generated simulated data: DJ. Produced figures: FC, MR and GY. Interpreted results: FC, MR, DJ and GY.

---

**Supplementary Figure 1. *DOT6* allele-replacement experiment.** Strains GY1566 (BY), GY1567 (RM) and GY1604, GY1605, GY1606, GY1607 (BY-*DOT6^RM^*) were cultivated in raffinose 2% and were shifted to a medium containing Raffinose 2% and Galactose 0.5%. After the indicated time, cultures were fixed with paraformaldehyde and analysed by flow cytometry. Histograms correspond to the fluorescent values obtained on cells gated for cell-size (see methods).

